# The ecological niche and seasonality of *Theileria equi* in Portugal

**DOI:** 10.64898/2026.11.02.763573

**Authors:** Ana Cabete, Jacinto Gomes, Ludovina Padre, Elisa Bettencourt, José Lourenço

**Affiliations:** MED - Mediterranean Institute for Agriculture, Environment and Development & CHANGE—Global Change and Sustainability Institute, Institute for Advanced Studies and Research, Universidade de Évora, Pólo da Mitra, Évora, Portugal; Department of Veterinary Medicine, School of Sciences and Technology University of Évora, Évora, Portugal; Elvas School of Biosciences, Polytechnic University of Portalegre, Portalegre, Portugal; VALORIZA - Research Centre for Endogenous Resources Valorisation, Polytechnic University of Portalegre, Portalegre, Portugal; CIISA - Centre for Interdisciplinary Research in Animal Health, Faculty of Veterinary Medicine, University of Lisbon, Avenida da Universidade Técnica, Lisbon, Portugal; Associate Laboratory for Animal and Veterinary Sciences (AL4AnimalS), Lisbon, Portugal; Universidade Católica Portuguesa, Católica Medical School, Católica Biomedical Research Centre, Portugal; Climate amplified diseases and epidemics (CLIMADE) consortium Europe, Portugal

## Abstract

Equine piroplasmosis, caused by the tick-borne protozoan *Theileria equi*, is an evolving concern in Europe. In recent years, it has been associated with substantial economic costs and with transmission showing signs of pushing into previously free or marginal northern regions. While piroplasmosis is endemic in Portugal, not much is known about the interplay of seasonally varying and spatial abiotic drivers that may modulate infection. In this study, we leveraged unique field-surveillance data based on equine PCR testing together with publicly available data to unravel and characterize the spatio-temporal dynamics and ecological niche of *Theileria equi*. Infection probability was estimated to increase with local bovine abundance, temperature and the presence of planosol soils, and to decrease with local equine density due to an intraspecific dilution effect of vector exposure. At an individual level, the highest infection probabilities were estimated among female and pasture-kept equids. We estimate strong seasonal signatures in local and national infection probability, maximized in August and minimized in February, and a spatially heterogeneous ecological niche landscape. Far northern and north-eastern regions combined low infection probability both in- and off-season with prolonged off-seasons, whereas centre and north-western regions combined high probability both in- and off-season with brief off-seasons. The former plausibly function as unsuitable reservoirs, and the latter as suitable year-round reservoirs. Our results provide a first quantitative, risk-stratified basis for regional and individual targeted *Theileria equi* surveillance, awareness and control in Portugal.

## Background

Equine piroplasmosis (EP) is an equid tick-borne disease. The first report of EP occurred in 1901, in South Africa [1,2]. By that time, the causative agent was named *Piroplasma equi*, later reclassified as *Babesia equi*. Years later it was discovered that two distinct parasites were capable of infecting equid erythrocytes. One of these was larger than the other, hence described as a “classic babesia”, corresponding to *Babesia caballi*. The smaller one, based on its size and the discovery of an extra-erythrocytic stage on lymphocytes, was considered as a “theileria-type” protozoan, being reclassified in 1998 and up to these days as *Theileria equi* [3]. Both hemoprotozoans belong to the Apicomplexa phylum, are globally distributed, but with a higher prevalence of *T. equi* when compared to *B. caballi* [4,5]. Mixed infections can occur but are less common. In endemic countries, most infected horses are asymptomatic, serving as reservoirs for tick infection and transmission of the disease [1]. In the case of *B. caballi*, the tick itself serves also as a reservoir, since transovarian infection is possible [1,6].

The importance of EP relates not only to animal welfare, in which clinical disease is typically characterized by fever, jaundice, anorexia, weight loss and poor performance (these last two specially in subclinical disease) [1,6], but also to the economic impact on the equine industry [4,7]. Veterinary treatment costs and resting equine athletes as a consequence of infection are two relevant economic factors, but there is also an impact on equine trade. Although objective estimates of the economic impact of EP are scarce, a 2009 outbreak in the USA generated approximately US$3 million in direct costs to a single equine operation, excluding lost market opportunities due to export restrictions [8]. In Spain, a study revealed that between 2015 and 2018, almost a quarter of the tested equines could not be exported to EP-free countries because of their seropositivity [7]. Currently, disease-free countries do not allow the entrance of EP-positive animals. Import requirements related to EP differ based on the classification of the country of both origin and destination [9]. In some countries, such as Australia [10], Canada [11], New Zealand [12], Japan [13] and the USA [14], official import-health requirements include pre-export and/or post-entry testing for *T. equi* and *B. caballi*. In Europe, although routine horse movements within the EU do not require proof of freedom from EP (Regulation (EU) 2016/429 [15]), non-endemic Member States can still implement national safeguards. An example is Ireland, where EP remains a notifiable disease under law (S.I. No. 130/2016) due to its historically disease-free status.

EP is known to have been present in Western Europe since the early 20th century [16]. In 1990, Friedhoff et al. [4], stated that “equine babesiosis extends from Portugal and Spain, through France and Italy”. To date, these countries remain endemic for EP [17–20]. Endemicity depends on a multitude of factors, not only on the parasite-vector-host cycle, but also on the environmental setting (e.g. climate, landscape, land use, etc), and the movement and management of animal hosts of both the hemoprotozoans and the ticks [21–23]. The free and frequent movement of equines throughout European countries, for example, enables the introduction and dissemination of EP in Europe, especially because asymptomatic horses serve as reservoirs. Tick abundance and its distribution, on the other hand, is known to be associated with climate [24,25], vegetation type [26,27], landscape [28,29], and the abundance and/or distribution of other animals besides equines [26,29]. Ultimately, the occurrence of EP is dependent on vector presence, with several ixodid ticks belonging to the genera *Dermacentor, Hyalomma* and *Rhipicephalus* recognised as EP vectors [1,6,30].

The related economic costs, impact on animal welfare and the link between multiple environmental factors and ticks, highlights the relevance of better understanding the ecological niche of hemoprotozoans causing EP. Only with such better understanding, can stakeholders work towards One Health insights that may be translatable for intervention, awareness and/or control [26,31] This is particularly the case for Europe, where equine movement is facilitated by intra-EU rules and climate change is on average accelerating more than in other continents [32,33].

In Portugal, the study of tick-borne pathogens dates back to the early 20th century. The fundamental lectures and reports by Professor Manuel Vianna, alongside the subsequent work of João Fraga de Azevedo in the mid-1930s [34,35], provided the earliest systematic mappings of tick vectors (*Rhipicephalus* and *Hyalomma spp.*) and early assessments of *Babesia* prevalence in the country. In the 1990s, Sabino Carvalho & Carvalho-Varela [36] reported the EP prevalence within the national equine population to be between 45 to 65%, a substantial baseline that established Portugal as endemic. Regional studies in mainland south [37] and north [38], as well as in the archipelago of the Azores [39] have reported the presence of infection. Recently, a retrospective study confirmed the endemicity in mainland Portugal [17]. Although EP is listed as a notifiable disease in the country, it remains notably underreported. Due to its endemic character, clinicians manage cases as part of their standard, daily routine, acting towards therapeutic intervention rather than systematically reporting.

Portugal’s vulnerability to climate impacts has been recognised in policy circles, with mean and extreme temperatures increasing over the past decades and projected to continue to do so [40]. The expected higher frequency and magnitude of drought periods may modify tick population dynamics, favouring the species adapted to a dyer Mediterranean environment, such as *Hyalomma lusitanicum*, *Hyalomma marginatum*, *Rhipicephalus bursa* and *Rhipicephalus annulatus*, all well established in the country [41,42]. Of the known factors that are understood to modulate EP distribution and occurrence, there remains a dearth of research related to the topic in Portugal, in particular regarding the interplay between abiotic seasonal and spatial drivers. At the same time, while there is potential in using modelling, it has so far not been applied to the ecology, prevalence, infection or seasonality of *T.equi* in Portugal.

In the current study, we have modelled the ecological niche of *T. equi* in Portugal, informed by a unique field-surveillance dataset and a range of available environmental and individual-host variables. We aimed for four objectives: to (1) characterize *T. equi’s* ecological niche(s) in the country; to (2) identify individual-host factors associated with *T. equi* in the country; to (3) estimate and characterize the national and regional seasonality of *T. equi*; and to (4) develop a first-of-a-kind regional category-based framework to inform near-future *T. equi* surveillance, awareness and control. We find that the ecological niche of *T. equi* in Portugal is driven by a mixture of environmental factors related to the distribution of specific mammal species, soil type and climate. Seasonality is spring-summer driven, with large differences between regions, not in timing but in absolute probability of *T. equi* infection. High risk regions for *T. equi* infection were found to be situated along a focal mid-center axis of the country, with an extra focal point in the littoral north, with both focal points including the largest metropolitan areas of Portugal - Porto and Lisboa. In parallel to environmental drivers, female equids and those living in pasture/field were found to be at higher risk of *T. equi* infection in the country.

## Methods

### Modelling data sources

#### Field study on T. equi infection

Data on *T. equi* infection was generated in a cross-sectional epidemiological study of EP in Portugal (unpublished). The study was designed to provide a geographically stratified coverage of the national equine population at the level of NUTS 3 (Nomenclature of Territorial Units). Each sample was accompanied by a questionnaire on sociodemographic and management factors. Only samples collected in continental Portugal (i.e. islands were not included) were considered for the present study, leaving 581 equid samples to analyse.

#### Sampling and diagnosis

Sampling was conducted from September 2023 until March 2026 by equine veterinarians working in the field and at an Equine Teaching Hospital (Faculty of Veterinary Medicine, University of Lisbon). The blood samples were collected from the jugular vein to an EDTA tube and kept refrigerated at 4°C until extraction. For this procedure, a commercial kit was used (Nucleic Acid Extraction Kit GF-1, Vivantis, Malaysia) and the obtained DNA stored at -20°C until molecular testing.

Samples were tested by PCR using a “catch-all” set of primers targeting the V4 hypervariable region of the 18S rRNA gene. The PCR reactions were carried out as described by Nagore et al. [43]. Positive samples were submitted to a second multiplex PCR, according to Alhassan et al. [44], to confirm which parasite was present and check for the presence of mixed infections.

#### Sociodemographic and management variables

The following variables were considered: NUTS 3, sex, age, breed and housing. Sex was a binary variable, with the male (M) and female (F) outcomes. Regarding age, it was divided into four categories: =<5, 5-10, >10-20; >20 years-old. Housing was also defined into four categories: box, paddock, pasture/field or mixed. As for breed, considering the diversity of answers in the questionnaires, it was divided as follows: Puro-sangue Lusitano (PSL; Purebred Lusitano), crossbreds, thoroughbreds, warmblood, pony, donkeys/mules and other/unknown.

#### Climate data sources

Climate data was obtained from Copernicus.eu “Essential climate variables for assessment of climate variability from 1979 to present”. Temperature, humidity and precipitation were averaged per NUTS 3 for the exact month-year of sampling records.

#### Land and vegetation data sources

Vegetation data was obtained from Copernicus.eu “Land cover classification gridded maps from 1992 to present derived from satellite observations” for the year 2019. We aggregated original categories into three overarching vegetation classes: natural (categories: 60, 70, 90, 100, 110, 120, 122, 130, 180), sparse (categories: 150, 151, 152, 153), trees (categories: 50, 60-62, 70-72, 80-82, 90, 100, 110). These three categories were then summarized as the mean proportion of land (that was *natural*, *sparse* or *trees*) within each NUTS 3.

#### Soil data sources

Soil type data was obtained from ISRIC - World Soil Information. Vertisols (clay-rich soils characterised by shrink–swell properties) and Planosols (soils with an abrupt textural change that may restrict drainage and promote seasonal waterlogging) were considered, summarized as the mean proportion of land (that was either *vertisol* or *planosol*) within each NUTS 3.

#### Mammal distribution data sources

We considered the spatial distribution of several mammal species: equines, caprines, bovines, swines, ovines and rabbits. Data was obtained from Geraldes et al. [45]. The data consists of the mean number of animals between the 2009 and 2019 per *freguesia* (the smallest geographical scale), which was summarized as the average per NUTS 3.

#### Vector distribution data sources

We considered the possible use of Portuguese surveillance data on the distribution of *Hyalomma marginatum*, *Hyalomma lusitanicum* and *Rhipicephalus sanguineus* ticks [41]. However, these data are made publicly available only as presence-absence records per year (no month recorded) without sampling effort or exact location detail, making it uninformative for the modelling pipeline of this study.

#### Data variables used for modelling

Pre-modelling, environmental variables were filtered by a pair-wise Pearson correlation threshold of 0.7. For each pair of variables correlated above the threshold, the variable with higher correlation with *T. equi* presence was kept. In this process, from the candidate list of variables, soil_mean, equines_mean, caprines_mean and ovines_mean were rejected. Since *T. equi* is an obligatory parasite of equines, we nonetheless decided to keep the variable equines_mean. Possible issues of collinearity due to this decision were evaluated at a later stage during model selection (detailed below). The final set of environmental variables considered for modelling is presented in **Table 1**, while the demographic variables considered for modelling are in **Table 2**. Final datasets used in modelling are provided as **Supplementary Tables S1** (PCR data and environmental variables) and **S2** (PCR data and demographic variables).

**Table 1 -.** Environmental variables, both part of the field-study data and public repositories used in modelling.

| variable | definition | source | ref. |
|---|---|---|---|
| MONTH | sampling month: 1 to 12 | this study | -- |
| SEASON | sampling season: spring, summer, autumn, winter | this study | -- |
| temp_mean | mean NUTS 3, monthly temperature in Celsius | Copernicus.eu | [46] |
| humi_mean | mean NUTS 3, monthly relative humidity in percent | Copernicus.eu | [46] |
| prec_mean | mean NUTS 3, monthly cumulative precipitation in mm | Copernicus.eu | [46] |
| natural_prop | mean NUTS 3, proportion of land that is natural vegetation | Copernicus.eu | [47] |
| sparse_prop | mean NUTS 3, proportion of land that is sparse vegetation | Copernicus.eu | [47] |
| trees_prop | mean NUTS 3, proportion of land that is tree cover | Copernicus.eu | [47] |
| planosol_prop | mean NUTS 3, probability of soil type being planosol | ISRIC - World Soil Information | [48] |
| vertisols_prop | mean NUTS 3, probability of soil type being vertisol | ISRIC - World Soil Information | [48] |
| bovines_mean | mean NUTS 3, number of bovines | literature | [45] |
| suines_mean | mean NUTS 3, number of suines | literature | [45] |
| rabbits_mean | mean NUTS 3, number of rabbits | literature | [45] |
| equines_mean | mean NUTS 3, number of equines | literature | [45] |

**Table 2 -.** Demographic variables, part of the field-study data used in modelling.

| variable name | definition |
|---|---|
| NUTS 3 | location of sampling |
| sex | sex of sampled horse |
| housing | housing status of sampled horse |
| age | age of sampled horse |
| breed | breed of sampled horse |

### Modelling pipeline

We used the binary outcome of PCR-testing each sampled animal as the probability (0-1) of detecting *T. equi* as a function of either the environmental or demographic variables detailed in the previous section. Continuous, numerical variables (i.e. environmental, excluding month and season) were standardized before modelling, to zero mean and unit variance [49]. We modelled the data using generalized linear models (GLMs) with binomial errors and a logit link [50] as detailed in the following sections.

#### Environment as determinant of *T. equi* infection

All variables were modelled as fixed effects, with the exception of MONTH and SEASON, which were modelled as random effects (categorical grouping variables). By default, for each animal sampled (sample unit) *i*, the probability of *T. equi* detection (herein termed infection probability) Π*_i_* was defined as (Model I): 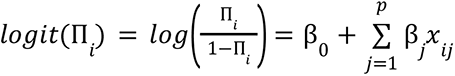, where β_0_ is an intercept, β*_j_* is a coefficient of the *j*-th predictor *x_ij_*, such that β*_j_* corresponds to the change in log-odds of *T. equi* detection Π_*i*_ per standard deviation change in the predictor. When considering the month and season of sample units as predictors, given that sample units within the same month and season cannot be fully considered as independent, a generalized linear mixed model (GLMM) with random intercepts was used instead [51], defined as 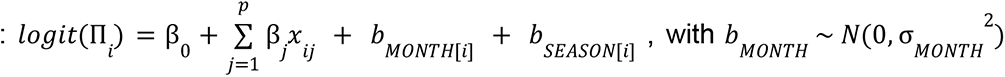 and 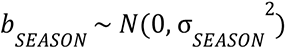

Introducing random intercepts allowed us to consider that the baseline log-odds of *T. equi* detection can shift per month or season, or both, thereby modelling temporal heterogeneity while keeping the fixed-effect structure parsimonious [52]. For modelling simplicity, under the context of limited input data from our field study, we did not consider NUTS 3 as a random effect, assuming that the environmental variables were already representative enough of the spatial dimension. Given this, four random effects structures were possible for modelling: no random effects (Model 1), random intercepts per month, per season, or both (Model 2).

With N=12 (Table 1) environmental variables and C=2 random effects as possible predictors, a large number of model formulations was possible. At a first stage, all possible model formulations were run under an automated model selection pipeline. All candidate models were ranked by the small-sample-corrected Akaike Information Criterion [53]. Since collinearity between predictors convey redundant information to modelling, multicollinearity was quantified for the predictors of each candidate model using the variance inflation factor (VIF)[54]. A strict heuristic was applied of rejecting any candidate model with at least one predictor presenting VIF ≥ 2.5 [55,56]. Of the remaining model candidates, the top 10 best-ranked AICc models were retained. This was applied for each of the four random effects structures, thus retaining a total of 40 candidate models (10 per random effect structure).

At a second stage, the short-listed 40 candidate models were evaluated in their predictive classification performance under a 5-fold cross-validation strategy. Threshold-independent performance was measured via operating characteristic curve (AUC) [57,58]. For threshold-dependent performance, a threshold was selected for each fold as the predicted-probability value (PPV) maximizing the sum of sensitivity and specificity ("Max Sens+Spec" criterion) [59,60]. Using this threshold, sensitivity (true positive rate), specificity (true negative rate), accuracy, true skill statistic (TSS) [61] and Cohen’s kappa (KAP) were quantified. Each of the metrics was then summarized as the mean ± standard deviation across the five held-out folds.

These two stages were designed to take as primary priority ecological interpretability and second priority predictive performance. The AICc / VIF strategy acted as primary selection criterion, and the myriad of performance metrics as secondary selection criterion towards selection of a single final model among the candidates. For this, we performed composite scoring. Performance metrics were min-max normalized (to 0 - 1), serving as input for the composite score formulation: 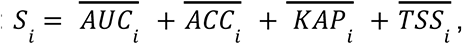, which was itself min-max normalized across model candidates. The model candidate with the highest composite score was selected as the final model.

#### In- and off-season score of relevance for *T. equi* infection

To summarise each NUTS 3 capacity for both infection intensity and persistence, we constructed a composite score from four local-level estimated quantities: the predicted probability of infection during the in-season (*P_in_*) and off-season (*P_off_*) periods, and the proportions of time spent in-season (*T_in_*) and off-season (*T_off_*). In- and off-season was determined by whether the estimated infection probability was above (in-season) or below (off-season) the predicted-probability value (PPV) maximizing the sum of sensitivity and specificity of the model. For this, we first computed a time duration-weighted infection intensity *i* = *P_in_* × *T_in_* + *P_off_* × *T_off_*. Because seasonal weights (*T_in_*, *T_off_* ) are considered as within NUTS 3 proportions, intensity *i* is scale-free and comparable across NUTS 3 regardless of differences in the number of sampling time points. We next computed an infection persistence ratio ρ = *min*(*P_off_* /*P_in_*, 1), defined as a dimensionless quantification of the proportion of in-season infection probability maintained during off-season. Infection intensity *i* and persistence ρ were then aggregated with a weighted geometric mean into score *R* = *i^a^* × ρ^1−*α*^ with *α* = 0. 5 as default. Geometric aggregation (and not additive aggregation) was used to limit compensability between intensity and persistence, e.g. a high intensity should not offset a complete loss of persistence potential. For mathematical integrity and to hold interpretation, NUTS 3 with *P_in_* = 0 were assigned ρ = 0, resulting in *R* = 0 (only one region, see Results). The score *R* was then normalized between 0 and 1 using the observed minimum and maximum to facilitate presentation.

#### Demographics as determinant of *T. equi* infection

Modelling of demographic variables followed the same general pipeline as modelling of environmental variables (running all candidate models, VIF assessment, AICc ranking, composite score selection). The exception was that all predictor variables (Table 2) were set to fixed effects and NUTS 3 as a random effect (without the consideration of random effects structures). A generalized linear mixed model (GLMM) with random intercepts was used [51], defined as (Model 3): 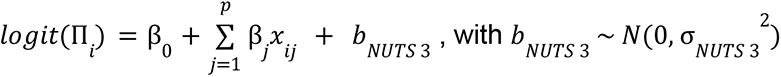

#### Computation

All work was performed in R (v4.6.0). GLMs were fitted with glm and mixed models with the package lme4 (v2.0.1). AICc computation was via the MuMIn package (v1.48.19). VIFs were computed with the car package (v3.1.5). AUC was calculated with the pROC package. Optimal thresholds and presence–absence evaluation with the PresenceAbsence package (v1.1.11). Data manipulation was performed with tidyverse (v2.0.0) and figures generated with the ggplot2 (v4.0.3) and cowplot (v1.2.0) packages.

## Results

### Field sampling

Horse blood samples were collected between September 2023 and March 2026 and registered by NUTS 3 geographical boundaries of Portugal. PCR testing effort varied with time and space (**Figure 1A-D**), sex (**Figure 1E**), housing (**Figure 1F**), breed (**Figure 1G**) and age (**Figure 1H**), presenting variation in terms of the proportions of positive and negative tests.

**Figure 1 -.**
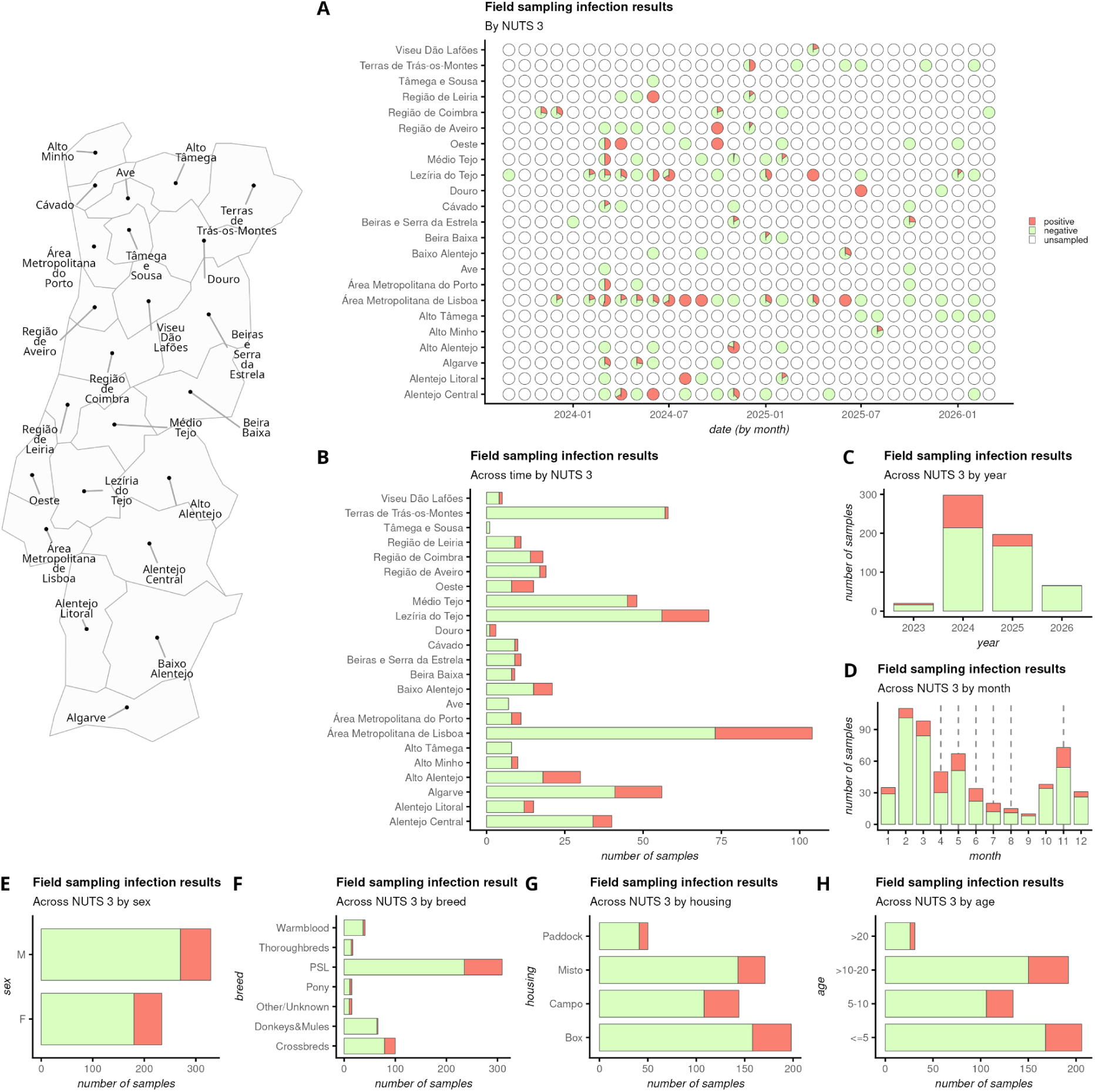
Field Sampling. The map on the left presents the NUTS 3 spatial boundaries of Portugal. (A) Results of T. equi PCR tests per date (year, month) by NUTS 3. Total number of samples (B) across the field-study time period by NUTS 3 according to PCR test result, (C) across all NUTS 3 by year, (D) across all NUTS 3 by month, (E) across NUTS 3 and time by sex, (F) across NUTS 3 and time by breed, (G) across NUTS 3 and time by housing, (H) across NUTS 3 and time by age-group. In panel D, the vertical dashed lines mark the months for which the proportion of positive PCR tests pass the 50% quantile of all months. Across panels A-H, PCR results are coloured by light green for negative, salmon for positive and white for unsampled (i.e. no PCR testing).

When aggregating across time, NUTS 3 regions that had *T. equi* infection (proportion of positive PCR) above the 50% quantile included Alentejo Litoral, Algarve, Alto Alentejo, Alto Minho, Baixo Alentejo, Douro, Lezíria do Tejo, Oeste, Região de Coimbra, Viseu Dão Lafões, Área Metropolitana de Lisboa and Área Metropolitana do Porto. Broadly, these regions were distributed across the country, excluding the far north.

NUTS 3 that were below the 25% quantile in the proportion of positive tests included Alto Tâmega, Ave, Cávado, Médio Tejo, Terras de Trás-os-Montes and Tâmega e Sousa (**Figure 1B**). All of these regions, with the exception of Médio Tejo situated in the mid-center of the country, were located in the far north. Considering only the years fully sampled from January to December, infection was higher in 2024 at ∼28%, than in 2025 at ∼15% (**Figure 1C**). When aggregated across years and NUTS 3, April to August and November were the months in which infection was above the 50% quantile of infection observed from January to December (**Figure1D**).

Regarding individual-host sex, across the entire field-study, ∼23% of female horses and ∼18% of males were positive (**Figure 1E**). Among samples with identified breed, infection was the highest at ∼27% for Ponies, ∼24% for Puro-sangue Lusitano (PSL; Purebred Lusitano), ∼21% for crossbreds, ∼18% for thoroughbreds, ∼10% for warmblood and ∼3% for donkeys/mules (**Figure 1F**). Horses living in pasture/field had an infection level of ∼25%, followed by those living in box at ∼20%, paddock with ∼18% and mixed with 16% (**Figure 1G**). Finally, mid-aged horses had the highest infection at ∼21% (same for both age-groups 5-10, >10-20), followed by the youngest 5-10 with ∼18% and the oldest >20 with ∼16% (**Figure 1H**). Overall, approximately 21% of all PCR tests performed were positive for *T. equi*.

### Environmental modelling

To explore, define and characterize the ecological niche of *T. equi* in Portugal, we modelled infection probability (PCR positivity) across samples as a function of environmental variables (**Table 1**; visually summarized in **Supplementary Figure 1**), following the pipeline described in the Methods section.

The selected model included the local number of bovines and equines, the proportion of planosol land and temperature as fixed effects, and the month of sampling as random effect (**Figure 2**). Model partial effects, where a particular variable is varied across its observed range and others are kept to their mean observed value, revealed that estimated infection probability of *T. equi* in Portugal is positively related to the number of bovines, planosol land and temperature, and negatively related with the number of equines (**Figure 2A**, **Supplementary Figure 2**).

**Figure 2 -.**
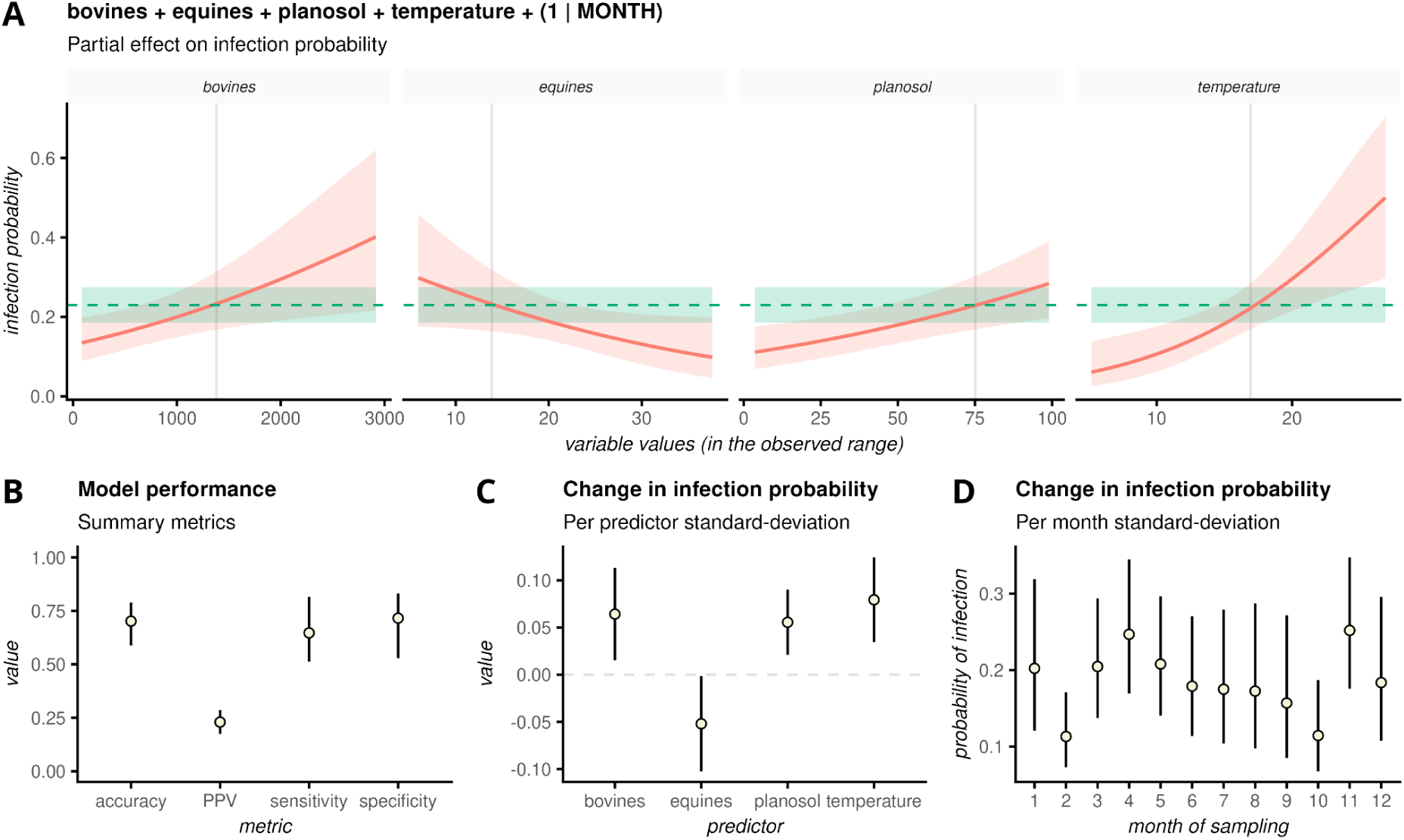
Selected environmental model. (A) Partial effects of fixed terms in the model, showing the variable specific relationship with the estimated infection probability of T. equi (red line: mean; red shade: standard error). The mean predicted-probability value (PPV) maximizing the sum of sensitivity and specificity of the model is presented by the shaded, green horizontal line, and its 95% CI by the green shaded area. The values of each variable (bovines, equines, planosol, temperature) above or below which the probability of T. equi is present breaches the PPV is marked by the vertical grey line on each subpanel. (B) Performance metrics for the model with mean (points) and 95% quantiles (whiskers). (C) Change in infection probability of T. equi presence per one-standard-deviation of each of the modelled variables (mean: points, 95% CI: whiskers). (D) Change in infection probability of T. equi presence per one-standard-deviation of sampling month (mean: points, 95% CI: whiskers).

Model performance was good (**Figure 1B**, **Supplementary Table S3**), with accuracy reaching 0.7 (95% CI, 0.59-0.79), specificity at 0.72 (0.53-0.83), and sensitivity equal to 0.65 (0.51-0.82) (**Figure 1D**). The predicted-probability value (PPV) maximizing the sum of sensitivity and specificity for the model was 0.23 (0.17-0.29).

From the partial effects, the infection probability was above the PPV when the number of bovines exceeded ∼1300, the number of equines was below ∼14, the proportion of planosol land was larger than ∼0.75, and temperature was warmer than ∼17 Celsius (**Figure 2A**). In effect, the infection probability of *T. equi* changed by +0.06 (standard error 0.03) per one-standard-deviation (one-SD) increase in the number of bovines, -0.05 (0.03) per one-SD increase in the number of equines, +0.06 (0.02) per one-SD increase in proportion of planosol land, and +0.08 (0.02) per one-SD increase in temperature (**Figure 1C**; all of which were statistically significant, i.e. p-value<0.05).

At the same time, since the model used sampling month as a random effect, it was possible to assess each month’s deviation in the baseline infection probability from that of an average month once environmental predictors had been accounted for (**Figure 2D**). In summary, infection probability changed by +0.07 (0-0.11) for a one-SD increase in the month-level random intercept (i.e. a month one-SD above the average month in baseline risk).

With the model fit to the field-study data, we estimated *T. equi* infection probability for years during which sampling was implemented (**Figure 3**). Monthly estimates revealed a strong seasonal signal and variation in the absolute levels of infection probability across NUTS 3 (**Figure 3A**). With one exception (Alto Tâmega), all NUTS 3 regions were seen crossing the mean predicted-probability value (PPV) for at least one month.

**Figure 3 -.**
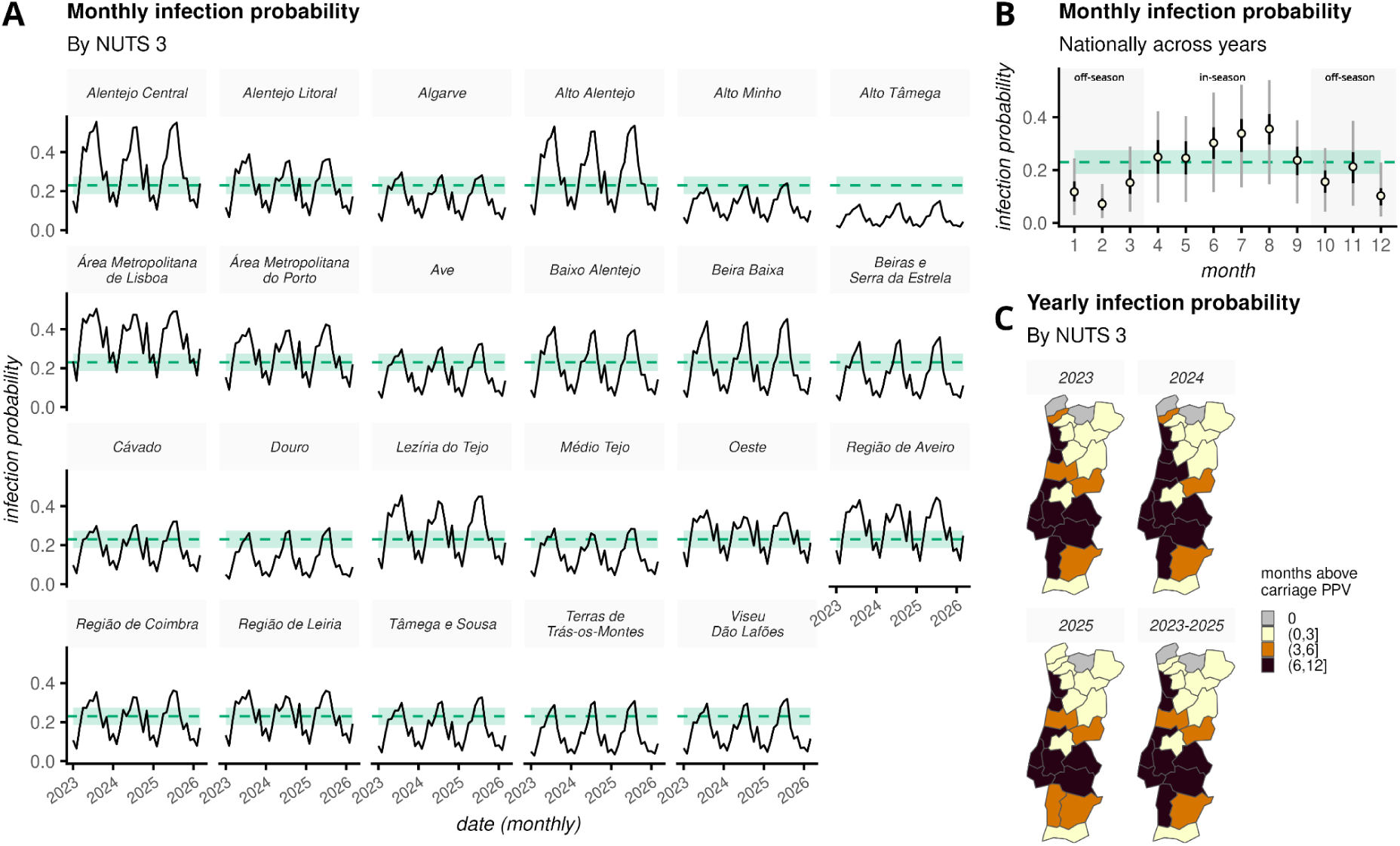
Seasonality of T. equi infection probability. (A) Estimated monthly T. equi infection probability (black line) during the field study time period by NUTS 3 (subpanel titles). The mean predicted-probability value (PPV) maximizing the sum of sensitivity and specificity of the model is presented by the shaded, green horizontal line, and its 95% CI by the green shaded area. (B) Estimated national, monthly T. equi infection probability when summarized across all years and NUTS 3 (points: mean; whiskers: black and grey ranges mark 50% and 95% quantiles, respectively). Year 2026 was excluded for not being complete. PPV estimates visualized as in panel A. (C) Yearly mean T. equi infection probability, by NUTS 3, per year and per entire time period of the field study. Year 2026 was excluded for not being complete. Color scale shows the ranges in number of months that estimated infection probability remained above the PPV (as per the estimations shown in panel A).

Nationally, i.e. when aggregated in space (NUTS 3) and time (years), monthly distributions of infection probability demonstrated a national seasonality signal characterized by lower probability in the winter months and by a gradual rise in probability during spring and summer (**Figure 3B**, **Supplementary Figure 3**). The national mean infection probability was estimated to be above the PPV between April and September, peaking in August (**Figure 3B**). These results were compatible with the seasonality observed in the proportion of positive PCR tests in the raw data (**Figure 1D**).

Under these seasonal patterns, in-season and off-season periods (probability above or below the PPV, respectively) could be defined as the month-ranges April-September and October-March, respectively (**Figure 3B**). Measuring seasonal signal as the absolute difference in probability between in-season and off-season periods, revealed that seasonal amplitude is stronger in the center and center-east regions of the country (**Supplementary Figure 4**).

When summarized per year by NUTS 3, infection probability remained above the PPV for at least 6 months in the central and northern-littoral regions of the country (**Figure 3C**). With the exception of two NUTS 3 (Médio Tejo in the center and Algarve in the extreme south), regions presenting 1 to 3 months in a year with infection probability below the PPV were situated in the extreme north and north-east of the country (**Figure 3C**). Regions presenting 0 months in a year with infection probability above the PPV (Alto Tâmega, Alto Minho) were all situated in the extreme north of the country (**Figure 3C**). These geographical patterns were broadly stable per year, between 2023 and 2025 (**Figure 3C**).

We next aimed at generating insights of relevance for surveillance, awareness and control from the estimated *T. equi* infection probability during in- and off-season periods. When analysing probability per NUTS 3 between 2023 and 2025, we found a positive correlation (0.83, p-value<0.05) between the proportion of time spent in-season and the mean probability levels during those time periods (**Figure 4A**). In contrast, we found a negative correlation (-0.75, p-value<0.05) when off-season (**Figure 4B**).

**Figure 4 -.**
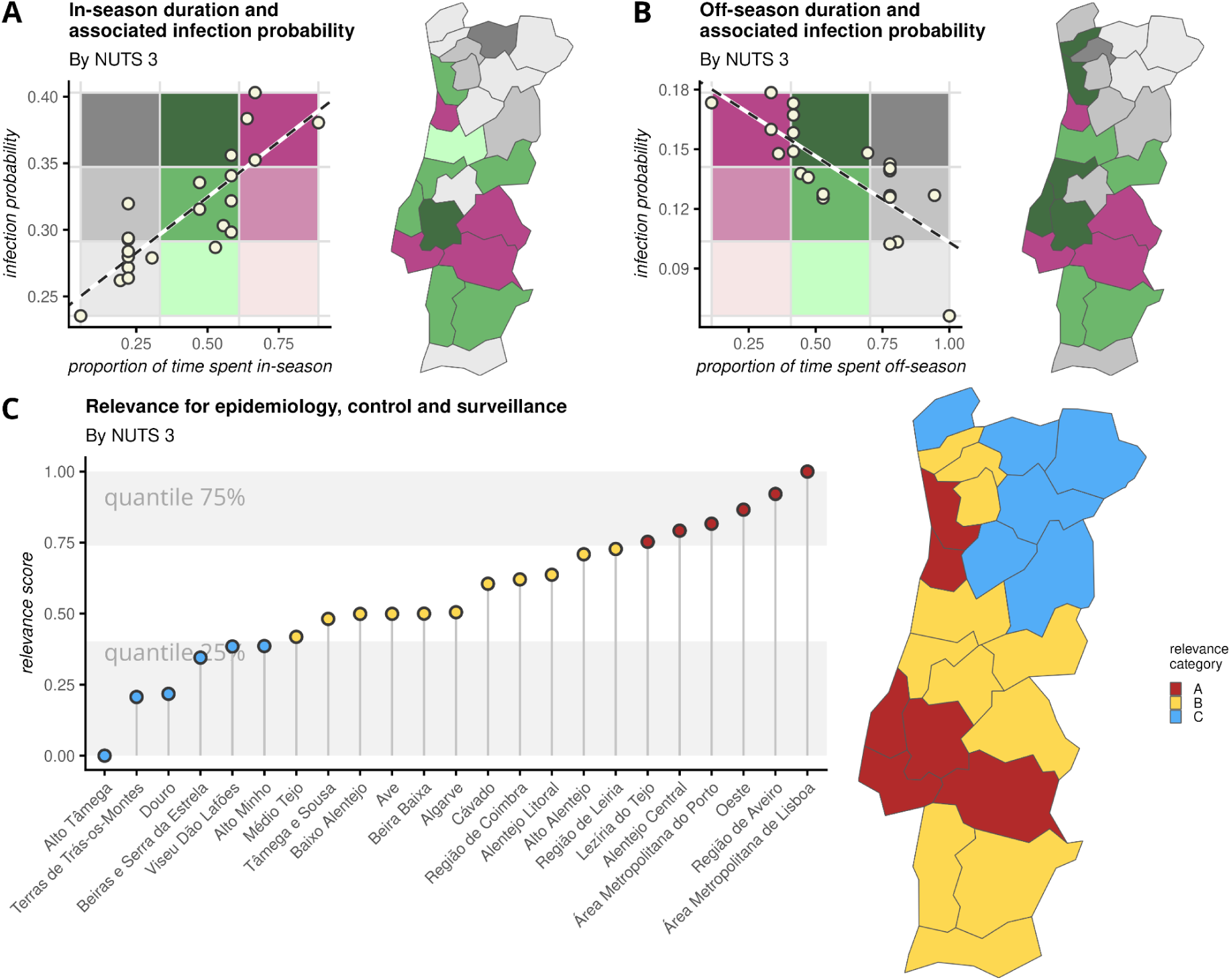
Patterns of relevance related to in- and off-season infection probability. (A) Relationship, per NUTS 3, of mean T. equi infection probability and time spent in-season (probability above PPV) for the period 2023-2025. (B) Same as panel A but for off-season periods (probability below PPV). (C) Relevance of both the relationships shown in panels A-B for epidemiology, control and surveillance of T. equi. Relevance is summarized into an aggregated score and further classified into 3 categorical levels A (highest relevance), B (intermediate relevance) and C (lowest relevance). The grey areas mark the quantile 75% (top) and 25% (bottom) of the relevance score range.

The subset of NUTS 3 that were in-season for the longest time periods with highest infection probability were also the ones being off-season for the shortest time periods with highest infection probability (**Figure 4A-B**), pointing to these regions playing a possible role in *T. equi* persistence throughout the year. These regions were located in the centre and centre-east, with the exception of a region in the littoral north (magenta, **Figure 4A-B**). In contrast, the subset of NUTS 3 that spent the least time in-season with the lowest infection probability, and were off-season for the longest time with lowest infection probability (shades of grey, **Figure 4A-B**), were all situated in the far north and north-east, with the exception of the Algarve in the far south. These patterns pointed to a possible weak role of such regions in persistence and transmission activity throughout the year.

We summarized the in- and off-season patterns (i.e. levels and time spent) of estimated *T. equi* infection probability (**Figure 4A-B**) into an aggregated score (see Methods, **Figure 4C**, **Supplementary Table S4**). The score was categorized as a 3-level framework by applying quantile thresholds: 0-25% (category C), 25-75% (category B) and 75-100% (category A). NUTS 3 categorized as class A represented regions with highest risk of *T. equi* infection throughout a typical year, having the highest infection probability both in- and off-season, and spending more time in-season than off-season. In contrast, NUTS 3 categorized as class C had the lowest infection probability both in- and off-season, and spent more time off-season than in-season. Consequently, those categorized as class B were regions presenting intermediate infection probability and balanced time spent in- and off-season. The highest risk regions with class A were identified in the mid-center axis of the country from the littoral to the interior and in a north-littoral focal point (**Figure 4C**). The lowest risk regions with class C were situated in the far north, in particular north-east of the country (**Figure 4C**).

### Demographic modelling

To explore and define individual-host factors that influence *T. equi* infection in Portugal, we modelled infection probability across samples as a function of demographic variables (**Table 2**; visually summarized in **Figure 1**), following the pipeline described in the Methods section.

The selected model included housing and sex of individual horses as fixed effects, and NUTS 3 as random effect (**Figure 5**). Model partial effects revealed that the strongest effects driving infection probability of *T. equi* were housing in pasture/field and female sex (**Figure 5A**). In contrast, the effect of housing in paddock and male sex were the weakest. Effectively, with reference to housing in box (odds=1), the odds-ratio was ∼1.36 (range 0.8-2.4) for pasture/field, ∼0.81 (0.46-1.45) for mixed and ∼0.78 (0.33-1.86) for paddock (**Figure 5B**). For sex, the likelihood of carrying *T. equi* was highest for females, with the odds-ratio of males being ∼0.7 (0.45-1.22) (**Figure 5B**). Of note, however, for both housing and sex, the odds-ratio upper or lower bounds crossed the reference level odds=1.

**Figure 5 -.**
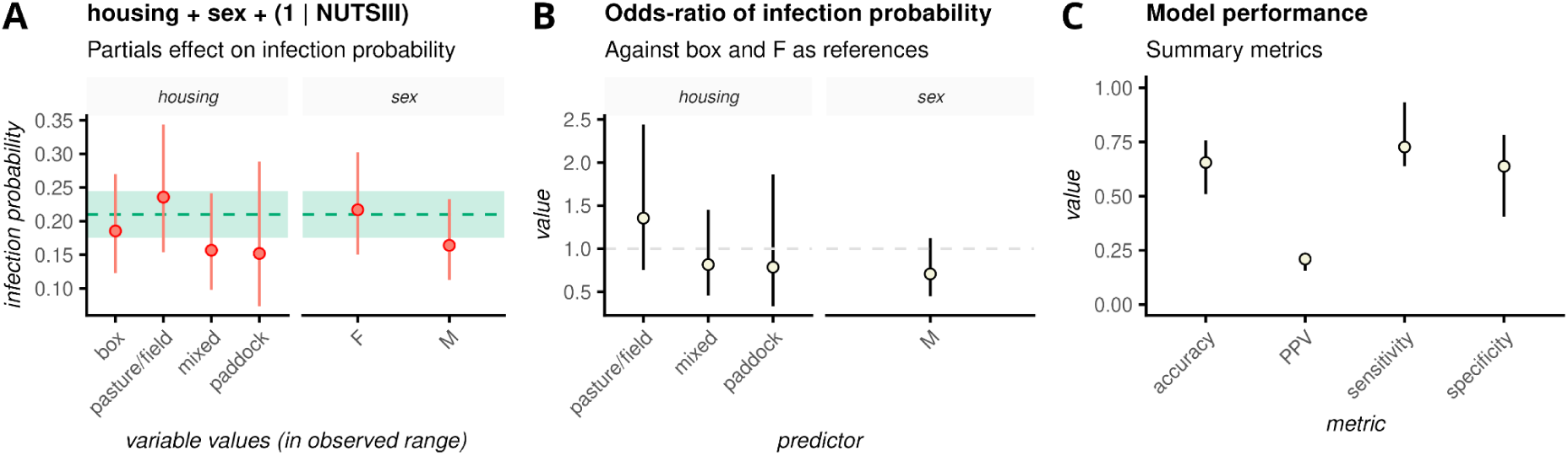
Selected demographic model. (A) Partial effects of fixed terms in the model, showing the variable specific relationship with the estimated infection probability of T. equi (red point: mean; red line: standard error). The mean predicted-probability value (PPV) maximizing the sum of sensitivity and specificity of the model is presented by the shaded, green horizontal line, and its 95% CI by the green shaded area. (B) Odds-ratio of T. equi infection probability (mean: points, 95% CI: whiskers). For housing, the reference value is box (i.e. box set to odds=1). For sex, the reference value is F (i.e. females set to odds=1). (C) Performance metrics for the model with mean (points) and 95% quantiles (whiskers).

Model performance was good (**Figure 5C**, **Supplementary Table S4**), with accuracy reaching ∼0.66 (CI 95%, 0.5-0.75), specificity ∼0.64 (0.4-0.8) and sensitivity ∼0.73 (0.64-0.93). The model’s predicted-probability value (PPV) maximizing the sum of sensitivity and specificity for the model was ∼0.21 (0.16-0.23).

## Discussion

In this study, we resorted to unique field-surveillance data based on PCR testing of equids for *T. equi*, together with publicly available environmental data, to investigate and characterize the ecological niche of the parasite in Portugal within a modelling framework.

We identified that its ecological niche is positively associated with the local number of bovines, proportion of planosol soil type and temperature, being also negatively associated with the number of local equines. In parallel to the determinants of ecological niche, female equines and those with pasture/field housing had the highest infection probability, while males and those with paddock housing had the lowest. The seasonality of estimated infection probability, both local and national, presented strong temporal signals. It typically increased during spring and summer with a peak in August, declining in autumn and winter with a through in February. Analysing estimated infection probability levels and time durations in- and off-season, revealed a clear national spatial structuring: regions in the far north and north-east had the least potential for *T. equi* positivity, while central west-to-east regions and a north-littoral region had the highest.

The recovered positive relationship between infection probability and temperature is well known and has a mechanistic basis [62]. Temperature is one of the major abiotic factors influencing tick ecology. For example, it modulates interstadial developmental rates [63], overwinter survival [64], the duration of host-seeking seasons [65,66] and parasite incubation periods [67,68]. In Portugal, tick species believed to be implicated in *T. equi* transmission (e.g. *Rhipicephalus bursa, Rhipicephalus annulatus, Dermacentor marginatus, Hyalomma marginatum and Hyalomma lusitanicum*) [26,42,69] are prosperous in central-southern half of the country, where the climate is classified as Warm Temperate Mediterranean [70] (being classified as Temperate Mediterranean in the north). Our results in terms of relevance score summarized as a 3-category framework, showing that the vast majority of regions with intermediate and highest relevance categories are concentrated in the center-south part of the country are consistent with such knowledge on the role of temperature.

In reference to tick species of relevance, cattle are known to be major maintenance hosts [26,42,71,72]. As such, regions with more bovines can, in principle, sustain denser tick populations. Equines sharing a habitat or housing conditions with cattle can thus be at higher risk of hosting ticks and thus of being infected with *T. equi*. This knowledge supports our results on the positive relationship between the local number of bovines and infection probability, in what can be described as an amplifying effect of cattle to tick populations.

The central role of moisture refugia in tick ecology is well known. Ticks can spend the vast majority of their life off-host, confined to the soil-litter interface where desiccation is a dominant source of mortality [73,74]. Concordingly, it has also been described that severe or long dry weather episodes depress tick abundance [75]. The planosol soil type is characterized as flat land with an impeded and slowly permeable subsoil that typically produces seasonal surface water stagnation, retaining moist well into local warmer and drier seasons. Land rich in planosol soil can thus act as a moisture reservoir in key times of the year, sustaining questing tick populations. Furthermore, the type of natural vegetation planosols typically support include the characteristic sparse grassland with scattered shrubs which are ideal grazing landscapes where animals including cattle, equines and other tick supporting wild-life can share pasture. The recovered planosol association with *T. equi* infection probability is thus supported by knowledge related to moisture-retentive, open grazing landscapes that link tick, wild and farmed animal population dynamics with the presence of the parasite.

In contrast, the negative association of the local number of equines with infection probability is compatible with the ecological concept of intraspecific dilution effect, in which an increase in the density of a host species (intraspecific variation) leads to a lower per-capita risk of infection [76,77]. Critical to this interpretation is that when equids are few, each horse may hold a large share of the local tick population, while when equids are abundant the tick population is divided among more horses, lowering each horse’s exposure to both the vector and the parasite. It may also be possible that husbandry intensity (e.g. contact with cattle, attendance to fairs, antiparasitic management, etc) is associated with the size of the horse population [18,78]. Larger populations may be better managed resulting in lesser tick infection and therefore parasite presence, while smaller populations may be less managed and/or associated with rough grazing. Discerning if this is the case in Portugal would require more detailed data, which we did not have access to at the time of the study.

The sex imbalance, expressed by the negative odds-ratio of male horses but with overlapping confidence to odds-ratio 1 (of females) is also supported by the published literature. A variety of reports find no significant association of *T. equi* infection with sex, while others, when reporting significant or marginal differences, find signals both in favour of males and females [23]. This literature incoherence, together with the overlapping odds-ratio signal we find, suggest that the marginal sex difference recovered from our model may be a proxy e.g. of animal management rather than a biological signal for which no discerning evidence exists in the literature. In Portugal, females are usually kept on pasture, specially the reproductive mares with their foals. This practice is believed to raise exposure to vectors, therefore increasing the probability of detectable parasitaemia. Our results are in agreement with previous studies that also associate sex imbalance with the management system rather than a biological factor itself [79].

Regarding housing settings, with reference to horses housed in box (i.e. stable), the higher odds-ratio of infection probability for pasture/field horses agrees with European reports finding that pasture access allows closer contact with wild-life and favourable tick-maintenance landscapes, favouring contact with questing ticks [80]. On the other hand, horses kept in box, besides being less exposed to ticks, are frequently groomed, drastically reducing the possibility of tick attachment and feeding. The lower odds-ratio for animals housed in mixed and paddock might reflect discrepancies in contact with ticks, livestock and wild-life habitats. Although these animals do have contact with the outdoors, in mixed systems this contact is usually temporary (i.e. during intermittent periods throughout the year). Regarding horses kept on paddocks, as they consist in smaller, limited areas that are kept clean, the risk is believed to be reduced. In both settings, horses receive higher attention and grooming than the ones kept only on pasture/field.

The positive correlation found between the estimated time spent in-season and mean in-season infection probability, suggest that regions with prolonged suitable seasons also experience higher infection probability. In contrast, the negative correlation found during off-season periods was suggestive that regions spending the longest time off-season also sustained the lowest infection probability during it. Intuitively, these patterns suggest that off-season periods in Portugal operate as an ecological bottleneck for *T. equi* transmission with duration and severity of the bottleneck varying geographically. Regions characterized by this behaviour were all found in the far north and north-east of Portugal. Similarly, regions combining long in-season periods and short off-season periods, both with highest infection probability (the centre axis and a focal north-littoral hotspot), therefore plausibly function as year-round reservoirs, favouring maintenance and infection in the equid population. Their location coincides with a Warm Temperate Mediterranean climate where tick species implicated in transmission may find suitable climate conditions, and where soil and host-population factors may maximize contact between equids, livestock, wildlife and questing ticks.

This study contains a number of data and methodological limitations of note. Infection was ascertained exclusively by PCR on blood samples. However, it is known that chronic *T. equi* carriers can show low and intermittent parasitemia, such that single-time-point PCR should be read as a lower bound of true infection. Equine field surveillance varied spatio-temporally, and consequently some of the analyses may partly reflect sampling design. Some of the results may also have had confounding by unmeasured management factors (e.g. husbandry intensity, antiparasitic practices, movement history, etc). Suitable tick distribution and density data was not available, such that some of the outputs could not be discerned in relation to direct spatio-temporal tick exposure landscapes. We decided to model environmental and demographic factors in separate frameworks, as it may provide insights to local stakeholders interested in distinct scales of action (e.g. interested or limited to considering environmental or demographic scales alone). Since it is likely that the considered factors are partly associated, their real-world effects share variance and should be read in the current study as complementary rather than independent or strictly additive. The study is relative to the 2023-2025 national scenario, which may not be fully representative of past and future scenarios in light of fast changing environmental conditions. In summary, the seasonal signatures and levels of infection probability and related quantitative framework presented in this study should be read as a unique and first-of-a-kind contribution, warranting reassessment as sampling accumulates, animal management data is better linked to PCR test results, publicly available tick surveillance data becomes more informative, and environmental conditions change.

In summary, using unique PCR field-surveillance data combined with publicly available environmental information from continental Portugal, we modelled the ecological niche of *T. equi*, estimating the spatio-temporal dynamics of infection probability. Estimated infection probability increased with local presence of bovines and planosol soils and with temperature, but decreased with the local number of equines. These signatures are consistent with cattle acting as amplifying maintenance hosts, temperature gradients shaping parasite and/or tick niches, moisture-retentive landscapes sustaining tick populations and/or favouring equine contact with infected ticks, and an intraspecific dilution of individual tick exposure with equine numbers. At the individual equine level only marginal differences were found, but were generally suggestive that infection probability was highest in female and free raging horses, compared to male and (partially-) housed animals. When analyzing spatio-temporal patterns of estimated infection probability between in- and off-season periods, a clear spatial structuring was identified. Of importance, regions along the central axis and a north-littoral hotspot showed strong in-season and off-season probability levels, suggesting their relevance for parasite maintenance and transmission. In the specific Portuguese setting, with focus on *T. equi,* our results provide novel insights into the environmental and demographic drivers of contemporaneous infection, and a first quantitative basis for near-future risk-stratified surveillance, awareness and control.

## Supplementary Files

**Tables S1** (Field-study PCR testing dataset with environmental variables), **S2** (Field-study PCR testing dataset with demographic variables), **S3** (Selected environmental model and summary measures of performance), **S4** (3-level framework categories) and **S5** (Selected demographic model and summary measures of performance) are provided as supplementary CSV files.

**Minimal R code** that implements the selected environmental model is provided as a supplementary R script.

## Acknowledgements

JL acknowledges the Fundação para a Ciência e Tecnologia (FCT) for institutional support to his host institution, CBR (DOI:10.54499/UID/06497/2025, UID/06497/2025).

## Data availability

Data is made available as supplementary tables.

## CRediT authorship contribution statement

ACC: Writing - review & editing, Writing - original draft, Validation, Investigation, Formal analysis, Data collection, Data curation, Conceptualization.

LP: Resources, Supervision, Funding Acquisition EB: Resources, Supervision, Funding Acquisition

JG: Writing - review & editing, Validation, Conceptualization, Supervision, Funding Acquisition, Project Administration.

JL: Writing - review & editing, Writing - original draft, Visualization, Validation, Software, Methodology, Investigation, Formal analysis, Data curation, Conceptualization.

## Supplemwntary Figures

**Supplementary Figure S1 -.**
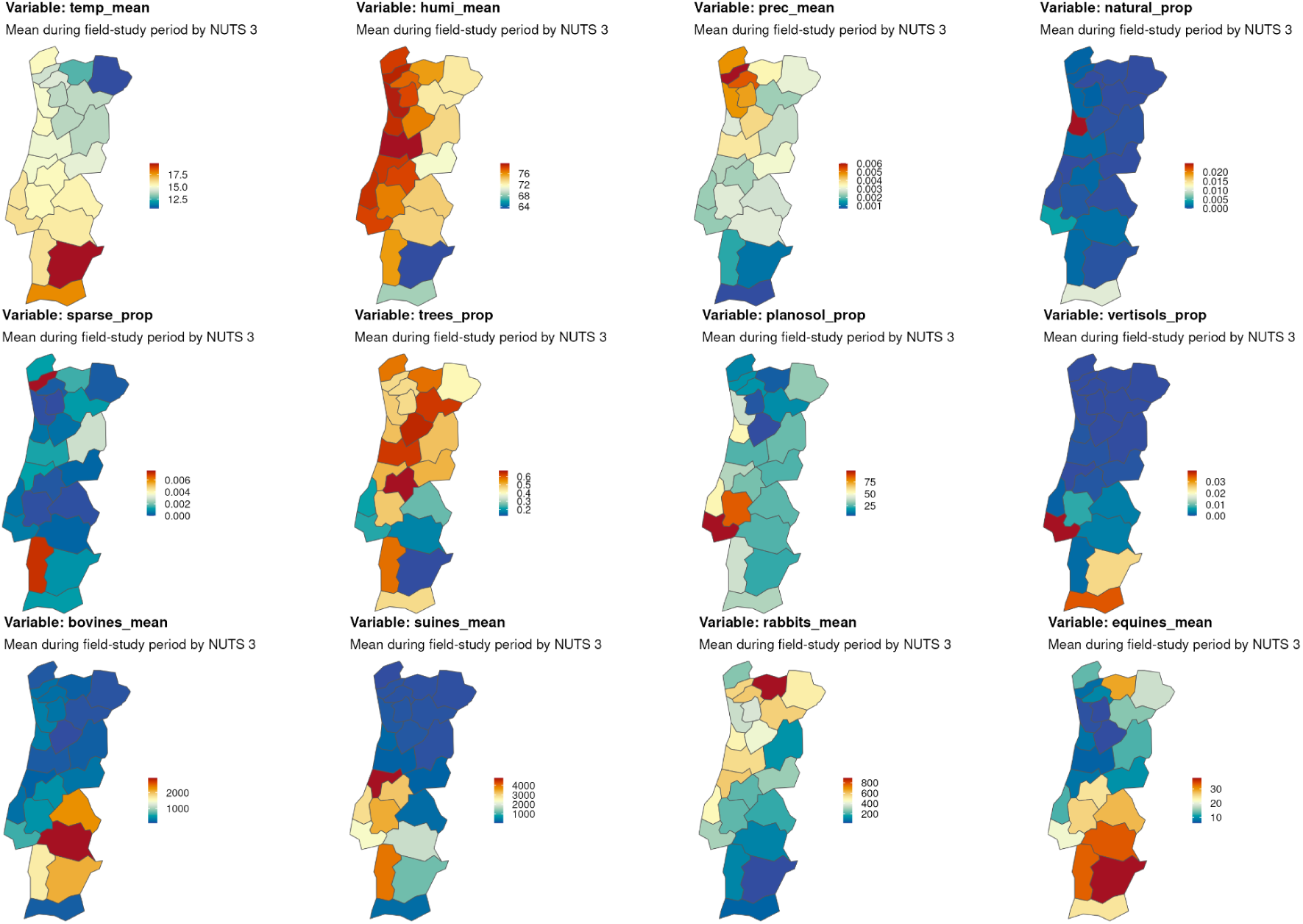
Summary of environmental variables used in modelling, visually presented by NUTS 3, averaged across the entire time period of the field-study.

**Supplementary Figure S2 -.**
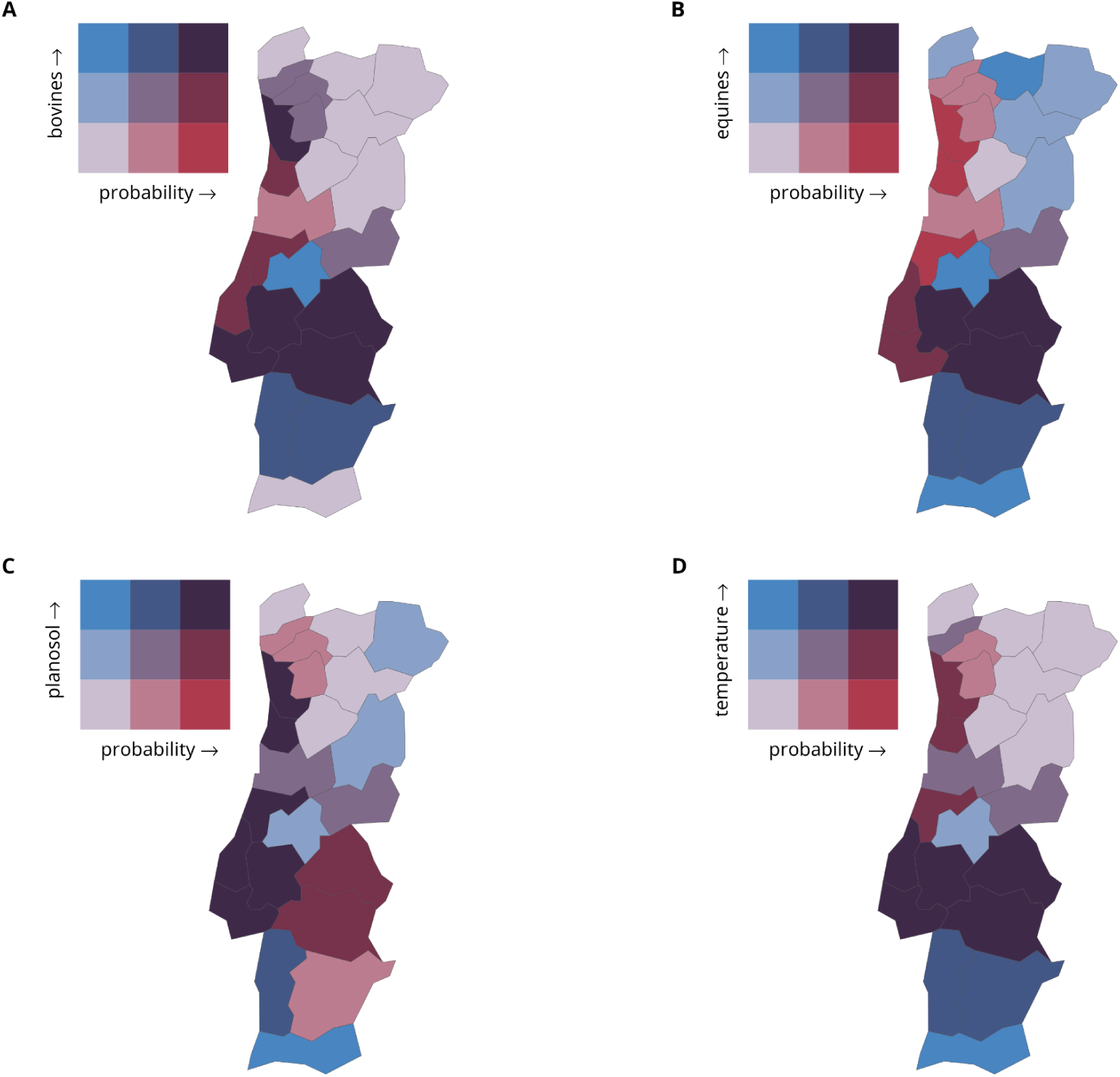
Summary of relationships between predictor variables and estimated carriage probability. Presented is the mean of predictor variables (A) bovines, (B) equines, (C) planosol and (D) temperature versus mean estimated carriage probability (coloured with the bivariate scheme) during the entire period of the field-study, per NUTS 3. The 3x3 bivariate scheme represents the combinations of each variable and carriage levels per 33% quantile of the values.

**Supplementary Figure S3 -.**
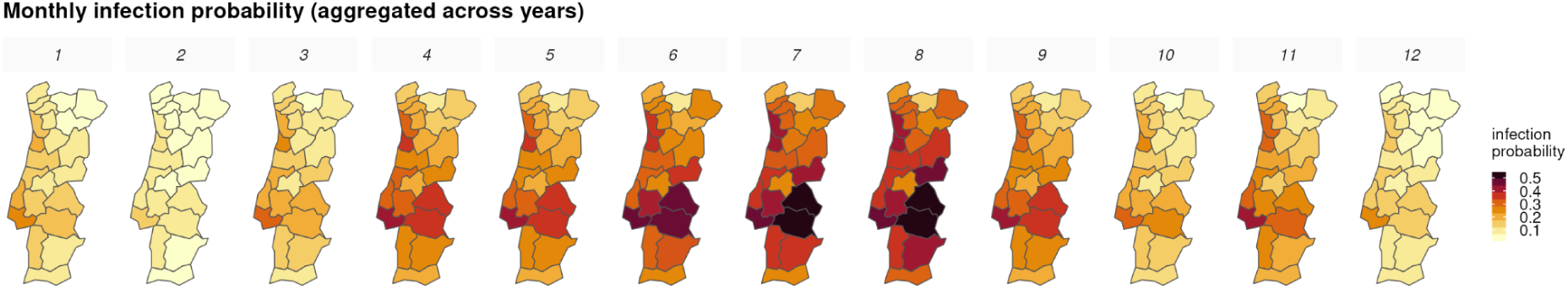
Summary of the seasonality of carriage probability. Presented per NUTS 3, estimated carriage probability is summarized per month (across all the years of the field-study period).

**Supplementary Figure S4 -.**
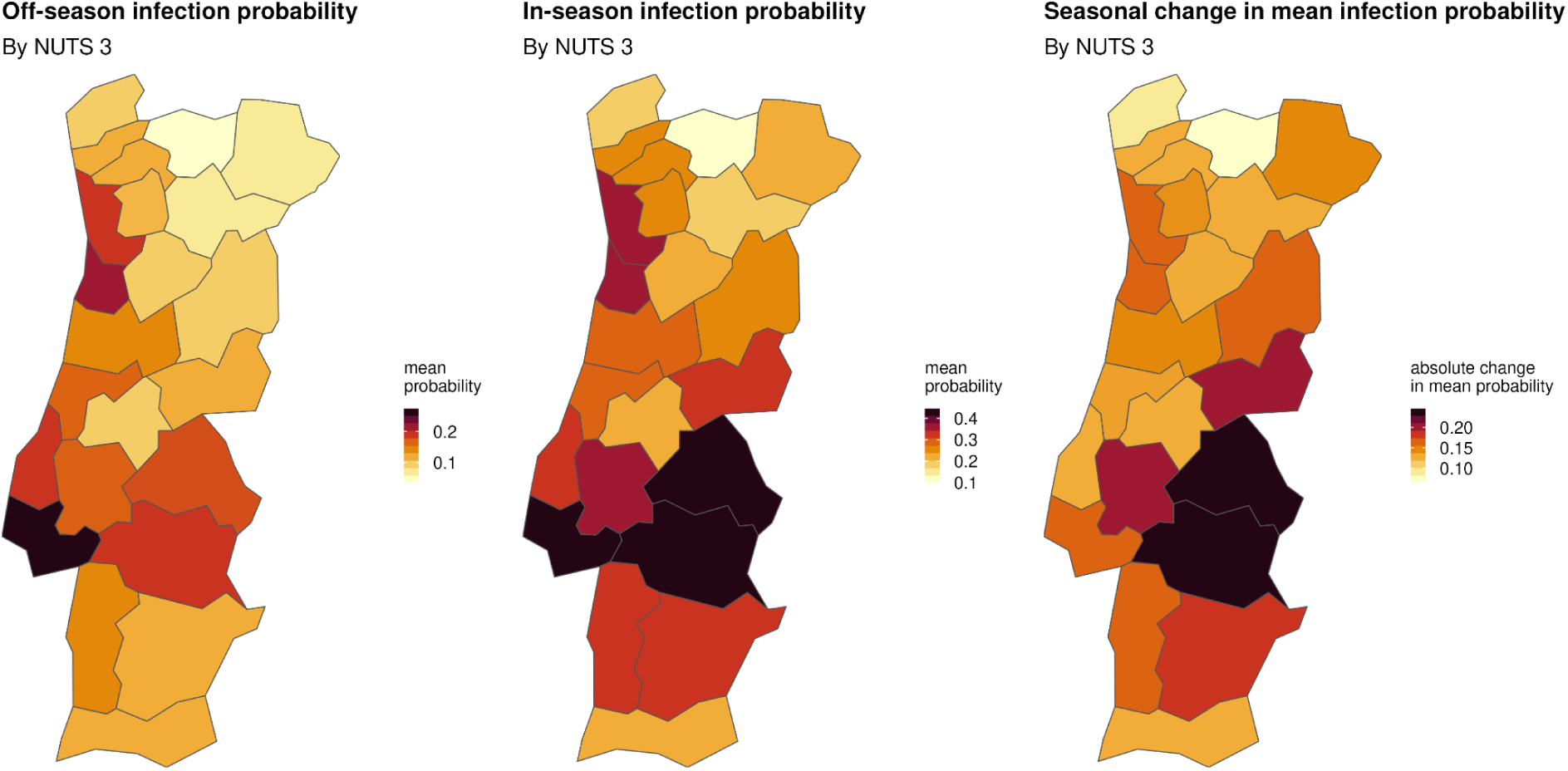
Summary of local seasonal signal. Off-season (left), in-season (middle) and the difference (right) between the two seasonal periods, in terms of estimated carriage probability, are presented by NUTS 3. Carriage probability is summarized by the mean for the entire period of the field study. Off- and in-season periods are determined as detailed in the main text. The difference between seasons (right) is calculated by the probability in-season minus the probability off-season.

